# Label-Free Prediction of Cell Painting from Brightfield Images

**DOI:** 10.1101/2021.11.05.467394

**Authors:** Jan Cross-Zamirski, Elizabeth Mouchet, Guy Williams, Carola-Bibiane Schönlieb, Riku Turkki, Yinhai Wang

## Abstract

Cell Painting is a high-content image-based assay which can reveal rich cellular morphology and is applied in drug discovery to predict bioactivity, assess toxicity and understand diverse mechanisms of action of chemical and genetic perturbations. In this study, we investigate label-free Cell Painting by predicting the five fluorescent Cell Painting channels from paired brightfield z-stacks using deep learning models. We train and validate the models with a dataset representing 1000s of pan-assay interference compounds sampled from 17 unique batches. The model predictions are evaluated using a test set from two additional batches, treated with compounds comprised from a publicly available phenotypic set. In addition to pixel-level evaluation, we process the label-free Cell Painting images with a segmentation-based feature-extraction pipeline to understand whether the generated images are useful in downstream analysis. The mean Pearson correlation coefficient (PCC) of the images across all five channels is 0.84. Without actually incorporating these features into the model training we achieved a mean correlation of 0.45 from the features extracted from the images. Additionally we identified 30 features which correlated greater than 0.8 to the ground truth. Toxicity analysis on the label-free Cell Painting resulted a sensitivity of 62.5% and specificity of 99.3% on images from unseen batches. Additionally, we provide a breakdown of the feature profiles by channel and feature type to understand the potential and limitation of the approach in morphological profiling. Our findings demonstrate that label-free Cell Painting has potential above the improved visualization of cellular components, and it can be used for downstream analysis. The findings also suggest that label-free Cell Painting could allow for repurposing the imaging channels for other non-generic fluorescent stains of more targeted biological interest, thus increasing the information content of the assay.

## 1 Introduction

Cell Painting [1] is a high-content assay used for image-based profiling in drug discovery [2]. It is an inexpensive methodology that captures the rich information in cell morphology and has shown promising utility in bioactivity prediction, identification of cytotoxicity and prediction of mechanisms of action [3] [4] [5]. Cell phenotypes are captured with six generic fluorescent dyes and imaged across five channels, visualizing eight cellular components: nucleus (DNA), endoplasmic reticulum (ER), nucleoli, cytoplasmic RNA (RNA), actin, Golgi, plasma membrane (AGP) and mitochondria (Mito) [1]. Promisingly, pharmaceutical companies such as Recursion Pharmaceuticals have implemented Cell Painting to support clinical stage pipelines [6].

Image-based assays suffer the technical limitation of having a finite number of imaging channels due to the requirement to avoid spectral overlap. Typically, there is a maximum of six stains which can be applied simultaneously across five imaging channels [7], thus hindering the ability to capture an even greater quantity of morphological information from other unstained subcellular compartments.

Advances in deep learning and computer vision have rapidly advanced image-based profiling, and is expected to accelerate drug discovery [2]. It is possible to use transmitted light images such as brightfield as an input to a convolutional neural network to generate the corresponding fluorescent images – so called *In Silico* labelling [8]. Brightfield imaging is ubiquitous in cell microscopy, and has been proposed as an informative imaging modality in itself. While brightfield imaging overcomes many drawbacks of fluorescent labelling, it lacks the specificity and clear separation of the structures of interest. Deep learning methods have shown promise in augmenting the information available in brightfield when trained with fluorescent signals as a domain transfer problem [9].

One way to achieve this is with conditional [10] generative adversarial networks (GANs) [11], introduced by Isola in 2016 (also known as pix2pix) [12]. From the input samples of an underlying unknown joint distribution of multi-modal data, the generator network can be trained to generate complementing paired data. The advantage of the conditional GAN is to overcome the limitations of a pixel-wise loss function such as L1 [13], used in many standard U-Net models.

In this study we investigate the utility of label free Cell Painting. Image-to-image approaches in Cell Painting are rare, with only known instance of GANs specifically for Cell Painting using Nucleoli, cytoplasmic RNA, Golgi plasma membrane and F-actin as an input to predict the other two fluorescent channels [14]. Fluorescent channels contain significantly more information than a brightfield z-stack, therefore predicting full fluorescent channels from brightfield images is a significantly more challenging task.

To the best of our knowledge this work is the first to demonstrate a label-free, five-channel Cell Painting replication from a transmitted light input modality such as brightfield. In this paper we have investigated the quality of the Cell Painting features extracted from the model-predicted image channels by correlating with the ground truth features. Additionally we attempt to replicate clustering by treatment group on a test sample of 273 fields of view from two different batches. This was achieved by training two models on a large, multi-batch dataset: a U-Net trained with L1 loss, and the same U-Net trained as the generator of a conditional Wasserstein [15] GAN. In order to fully interrogate the utility of the label-free Cell Painting image channels, a comparison of both image-level metrics and morphological feature predictions between the two models is presented, contextualizing the non-normalized image metrics [16] and the feature-level evaluation results. We have aimed to provide a greater understanding to what extent convolutional neural network-based approaches may be able to assist or replace fluorescent staining in future studies.

## 2 Materials and Methods

### 2.1 Cell Culture and Seeding

A U-2 OS cell line was sourced from AstraZeneca’s Global Cell Bank (ATCC Cat# HTB-96). Cells were maintained in continuous culture in McCoy’s 5A media (#26600023 Fisher Scientific, Loughborough, UK) containing 10% (v/v) fetal bovine serum (Fisher Scientific, #10270106) at 37°C, 5% (v/v) CO_2_, 95% humidity. At 80% confluency, cells were washed in PBS (Fisher Scientific, #10010056), detached from the flask using TrypLE Express (Fisher Scientific, #12604013) and resuspended in media. Cells were counted and a suspension prepared to achieve a seeding density of 1500 cells per well using a 40 μL dispense volume. Cell suspension was dispensed directly into assay-ready (compound-containing) CellCarrier-384 Ultra microplates (#6057300 Perkin Elmer, Waltham, MA, USA) using a Multidrop Combi (Fisher Scientific). Microplates were left at room temperature for 1h before transferring to a microplate incubator at 37°C, 5% (v/v) CO_2_, 95% humidity for a total incubation time of 48 hours.

### 2.2 Compound Treatment

All chemical compounds were sourced internally through the AstraZeneca Compound Management Group and prepared in stock solutions ranging 10-50mM in 100% DMSO. Compounds are transferred into CellCarrier-384 Ultra microplates using a Labcyte Echo 555T from Echo-qualified source plates (#LP-0200 Labcyte, High Wycombe, UK). Compounds were tested at multiple concentrations; either 8 concentration points at 3-fold (half log) intervals or 4 concentration points at 10-fold intervals (supplementary Table A contains concentration ranges for compound collections tested). Control wells situated on each plate consisted of neutral (0.1% DMSO v/v) and positive controls (63nM mitoxantrone, a known and clinically-used topoisomerase inhibitor and DNA intercalator). Compound addition to microplates was performed immediately prior to cell seeding to produce assay-ready plates.

### 2.3 Cell Staining

The Cell Painting protocol was applied according to the original method [1] with minor adjustments to stain concentrations. Hank’s balanced salt solution (HBSS) 10x was sourced from AstraZeneca’s media preparation department and diluted in dH2O and filtered with a 0.22μm filter. MitoTracker working stain was prepared in McCoy’s 5A medium. The remaining stains were prepared in 1% (w/v) bovine serum albumin (BSA) in 1x HBSS (Table A supplementary).

Post incubation with compound, media was evacuated from assay plates using a Blue®Washer centrifugal plate washer (BlueCatBio, Neudrossenfeld, Germany). 30μL of MitoTracker working solution was added and the plate incubated for a further 30 min at 37°C, 5% CO_2_, 95% humidity. Cells were fixed by adding 11μL of 12% (w/v) formaldehyde in PBS (to achieve final concentration of 3.2% v/v). Plates were incubated at room temperature for 20 min then washed using a Blue®Washer. 30μL of 0.1% (v/v) Triton X-100 in HBSS (#T8787 Sigma Aldrich, St. Louis, MO, USA) solution was dispensed and incubated for a further 20 min at room temperature followed by an additional wash. 15μL of mixed stain solution was dispensed, incubated for 30 min at room temperature then removed by washing. Plates were sealed and stored at 4°C prior to imaging.

### 2.4 Imaging

Microplates were imaged on a CellVoyager CV8000 (Yokogawa, Tokyo, Japan) using a 20x water-immersion objective lens (NA 1.0). Excitation and emission wavelengths are as follows for fluorescent channels: DNA (ex: 405nm, em: 445/45nm), ER (ex: 488nm, em: 525/50nm), RNA (ex: 488nm, em: 600/37nm), AGP (ex: 561nm, em: 600/37nm) and Mito (ex: 640nm, em: 676/29nm). The three brightfield slices are from different focal z-planes; within, 4μm above and 4μm below the focal plane. Images were saved as 16-bit .tif files without binning (1996×1996 pixels).

### 2.5 Pre-Processing

The selected images were bilinearly downscaled from 1996×1996 to 998×998 pixels to reduce computational overheads. Global intensity normalization was implemented to eliminate intensity differences between batches. For each channel, each 998×998 image was constrained to have a mean pixel value of zero and a standard deviation of one. Corrupted files or wells with missing fields were removed from the usable dataset and replaced with files from the corresponding batch.

### 2.6 CellProfiler Pipeline

A CellProfiler [17] [18] pipeline was utilized to extract image- and cell-level morphological features. The implementation followed the methodology of Way *et al.* 2021 [19], and is chosen as a representative application of Cell Painting. The pipeline we used is included in our GitHub repository^1^. CellProfiler was used to segment nuclei, cells and cytoplasm, then extract morphological features from each of the channels. Single cell measurements of fluorescence intensity, texture, granularity, density, location and various other features were calculated as feature vectors.

### 2.7 Morphological profile generation

Features were aggregated using the median value per image. For feature selection, we adopted the following approach:

- Drop missing features - features with > 5% NaN values, or zero values, across all images
- Drop blocklisted features [20] which have been recognized as noisy features or generally unreliable
- Drop features with greater than 90% Pearson correlation with other features
- Drop highly variable features (>15 SD in DMSO controls)

This process reduced the number of useable features to 611, comparable to other studies. We used the ground truth data only in the feature reduction pipeline to avoid introducing a model bias to the selected features.

### 2.8 Training and Test Set Generation

We sampled from 17 of the 19 batches to select wells for training (Table 1). The remaining two batches were used to select the test set, and were excluded in the training process. Compounds from the test set were comprised from a publicly available list of known pharmacologically active molecules, with a known observable phenotypic activity. We randomly sampled 3000 wells for model training and hyperparameter tuning, with the constraint to force an overall equal number of wells per batch. We randomly selected one field of view from each of the four fields in the well, which was the image used in the training set. For model tuning we used 90/10 splits sampled randomly from the training set, before using the full training set to train the final models. The test set contained 273 images and was chosen by sampling randomly within each treatment group across the two remaining batches (treatment breakdown in Table 1).

**Table 1:** The composition of the Training, Validation and Test sets. Choosing data from many batches and experiments should make our models learn more robust, common features, which gives us the best chance of successful prediction on unseen data.

|  | No. of Batches | No. of 384-well Plates | No. of Images | Controls and Treatment Distribution | Collection Description (no. of compounds) |
| --- | --- | --- | --- | --- | --- |
| **Total Available** | 19 | 140 | 150,000 (4 fields per well) | 2.1% positive controls; 6.4% negative controls; 91.5% treatments | Mixed collection containing a high chemical diversity set with a range of physicochemical properties ($\sim 7,000$); a phenotypic set ($\sim 1,000$) and a set of pan-assay interference compounds ($\sim 1,000$) |
| **Training / Validation Set** | 17 | 124 | 3,000 (1 field of view used per well) | Randomly sampled from above | Randomly sampled from above |
| **Test Set** | 2 | 16 | 273 (1 field of view used per well) | 26 positive controls; 77 negative controls; 170 treatments | Publicly available phenotypic set (exclusively in test set) |

### 2.9 U-Net with L1 Loss Network Architecture

Our first model is based on the original U-Net [21], a convolutional neural network which has proven very effective in many imaging tasks. U-Net architectures have been used to solve inverse reconstruction problems in cellular [22], medical [23] and general imaging problems [24]. For segmentation tasks, out of the box U-Net based architectures such as nnU-Net [25] have been proven to perform very well even compared to state of the art models.

U-Nets involve a number of convolutions in a contracting path with down sampling or pooling layers, and an expansive path with up sampling and concatenations, allowing for retention of spatial information while learning detailed feature relationships. The network captures multi-scale features of images through different resolutions by going up and down the branches of the network.

We adapted the typical grayscale or RGB channel U-Net model to have 3 input channels and 5 output channels to accommodate our data. An overview of the model network and training is presented in Figure 2. There were 6 convolutional blocks in the downsampling path, the first with 32 filters and the final with 1024 filters. Each block performed a 2d convolution with a kernel size of 3 and a stride of 1, followed by a ReLU then batch normalization operation. Between blocks a max pooling operation with a kernel size of 2 and a stride of 2 was applied for downsampling. The upsampling path was symmetric to the downsampling, with convolutions with a kernel of 2 and a stride of 2 applied for upsampling. The corresponding blocks in the contracting and expansive paths were concatenated as in the typical U-Net model. The final layer was a convolution with no activation or batch normalization. In total our network had 31×10^6^ trainable parameters.

**Figure 1:**
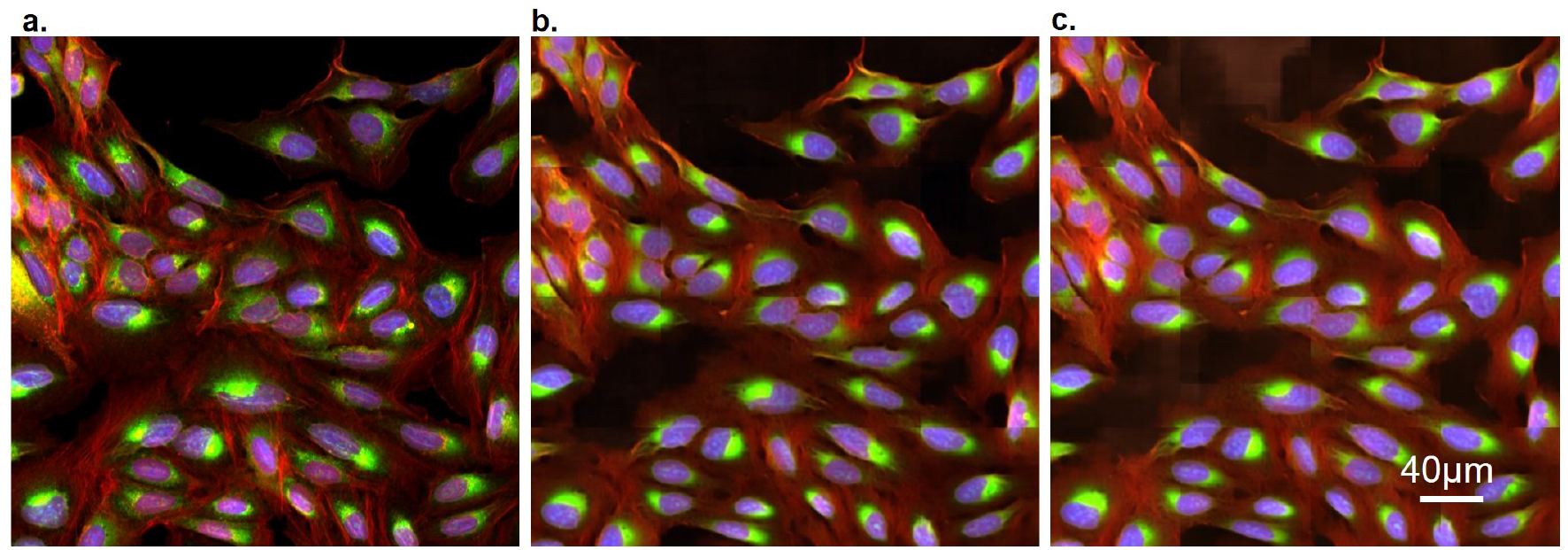
Colored composite image of three channels from the test dataset (Table 1): AGP (red), ER (green) and DNA (blue). a) Ground truth. b) cWGAN-GP model output. c) U-Net model output. Each image is 512×512 pixels.

**Figure 2:**
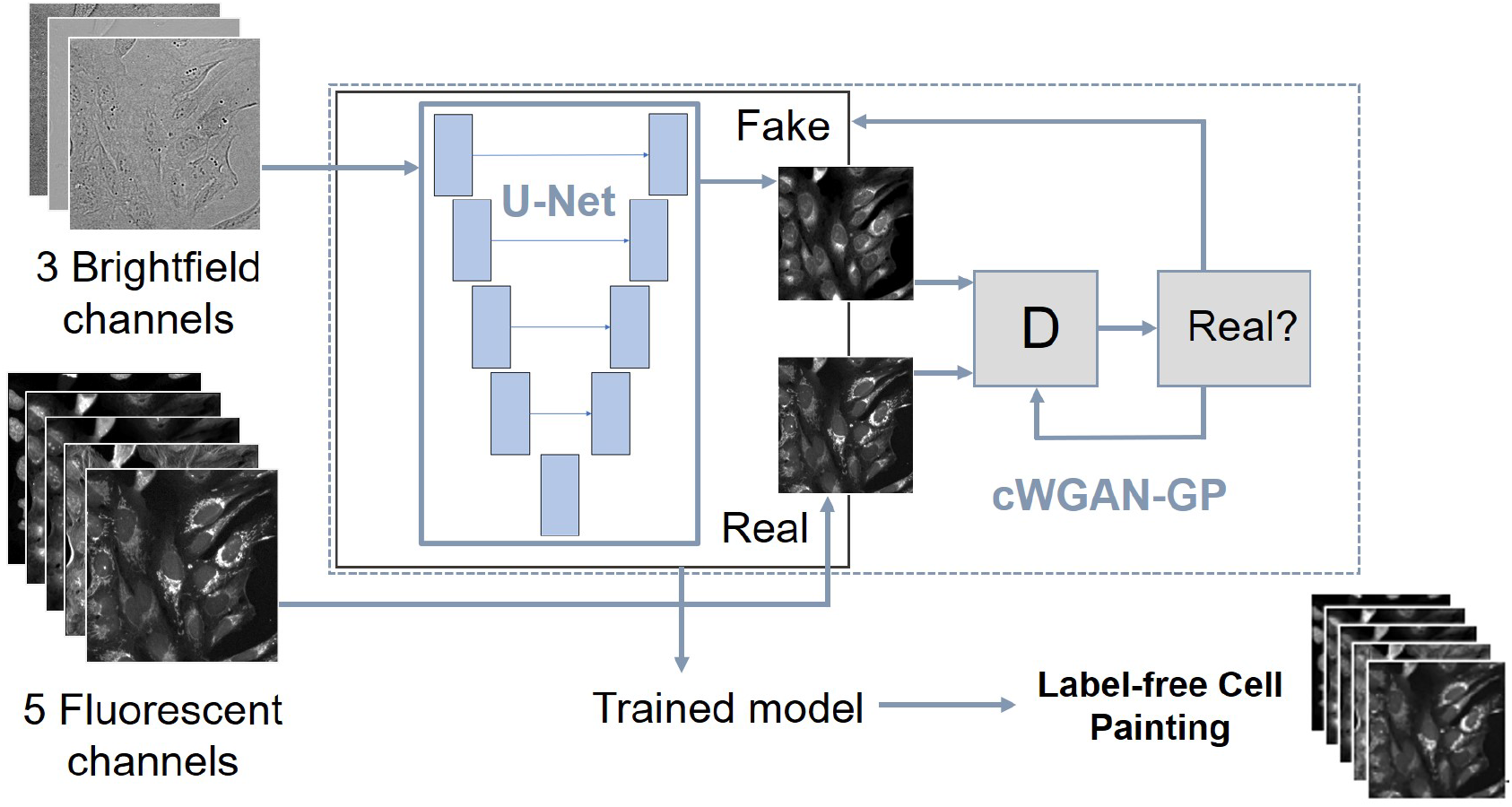
Summary of the models: U-Net trained on L1 loss and cWGAN-GP.

For pairs of corresponding images (*x*_*i*_, *y*_*i*_), where *x*_*i*_ is a 3 channel image from the brightfield input space and *y*_*i*_ is a 5 channel image from the real fluorescent output space, the loss function 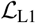 for the U-Net model is:

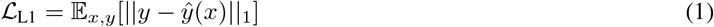

where 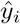 is the predicted output image from the network.

### 2.10 Additional Adversarial (GAN) Training Network Architecture

The second model we trained is a conditional [10] GAN [11], where the generator network *G* is the same U-Net used in the first model. Typically, GANs have two components: the generator *G* and the discriminator *D*. The generator and the discriminator are simultaneously trained neural networks: the generator outputs an image and the discriminator learns to determine if the image is real or fake. For a conditional GAN, the objective function 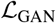 from a two-player minimax game is defined as:

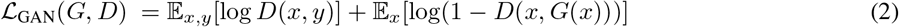

where *G* is trying to minimise this objective, and *D* is trying to maximize it

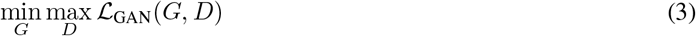

Many GAN image reconstruction models follow the framework from Isola [12] where the model objective function *G*\* is, for example, a mix of both objective functions:

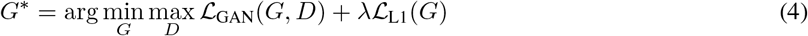

for some weighting parameter λ.

These constructions can be unstable and difficult to train, and this was the case for our dataset. To overcome difficulties with training we opted for a conditional Wasserstein GAN with gradient penalty [26] (cWGAN-GP) approach. This improved Wasserstein GAN was designed to stabilize training, useful for more complex architectures with many layers in the generator.

The Discriminator network *D* - alternatively the critic in the WGAN formulation - is a patch discriminator [12] with the concatenated brightfield and predicted Cell Painting channels as the eight-channel input. There were 64 filters in the final layer and there were three layers. For the convolutional operations the kernel size is 4 and the stride is 2. The output is the sum of the cross-entropy losses from each of the localized patches.

In equation (4), the L1 loss term enforces low-frequency structure, so using a discriminator which classifies smaller patches (and averages the responses) is helpful for capturing high frequency structures in the adversarial loss. For Wasserstein GANs the discriminator is called the critic as it is not trained to classify between real and fake, instead to learn a K-Lipschtiz function to minimize the Wasserstein loss between the real data and the generator output.

Hence, for our conditional WGAN-GP-based construction we trained the generator to minimize the following objective:

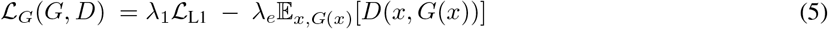

Where λ_1_ is a weighting parameter for the L1 objective. We introduce λ_e_ as an adaptive weighting parameter to prevent the unbounded adversarial loss overwhelming L1:

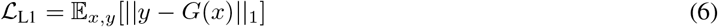

The critic objective, which the network is trained to maximize, is:

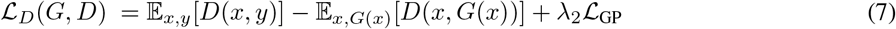

Where λ_2_ is a weighting parameter for the gradient penalty term:

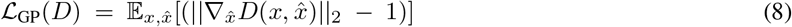

which is used to enforce the K-Lipschitz constraint. 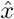 is from uniformly sampling along straight lines between pairs of points in the real data and generated data distributions [27].

### 2.11 Model Training and Computational Details

From the training set, random 256×256 patches were cropped to serve as inputs to the network. No data augmentation was used. The models were trained on the AstraZeneca Scientific Computing Platform with a maximum allocation of 500G memory and four CPU cores. The PyTorch implementation of the networks and training is given in our GitHub repository. The U-Net model was trained with a batch size of 10 for 15,000 iterations (50 epochs). The optimizer was Adam, with a learning rate of 2×10^−4^ and weight decay of 2×10^−4^. The total training time for the U-Net model was around 30 hours.

The cWGAN-GP model was trained with a batch size of 4 for an additional 21,000 iterations (28 epochs). The generator optimizer was Adam, with a learning rate of 2×10^−4^ and β_1_ of 0 and β_2_ of 0.9. The critic optimizer was Adam with a learning rate of 2×10^−4^ and β_1_ of 0 and β_2_ of 0.9 and a weight decay of 1×10^−3^. The generator was updated once for every 5 critic updates. The L1 weight is λ_1_ = 100 and the gradient penalty weighting parameter is λ_2_ = 10. λ_e_ = 1/epoch. The total training time for the cWGAN-GP model was an additional 35 hours. For each model, the best performing epoch was selected by plotting the image evaluation metrics of the validation split for each epoch during training. Once the metrics stopped improving (or got worse), training was stopped as the model was considered to be overfit.

### 2.12 Model inference

Input and output images were processed in 256×256 patches. The full-sized output images were stitched together with a stride of 128 pixels. This was chosen as half of the patch size so the edges met along the same line, which reduced the number of boundaries between tiles in the full-sized reconstructed image. Each pixel in the reconstructed prediction image for each fluorescent channel is the median value of the pixels in the four overlapping images. The full-sized, restitched images used for the metric and CellProfiler evaluations were 998×998 pixels for each channel, examples of which are displayed in supplementary Figure B.

### 2.13 Image-Level Evaluation Metrics

The predicted and target images were evaluated with five metrics: mean absolute error (MAE), mean squared error (MSE), structural similarity index measure (SSIM) [28] [29] peak signal-to-noise ratio (PSNR) [29] and the Pearson correlation coefficient (PCC) [8] [30]. MAE and MSE capture pixel-wise differences between the images, and low values are favorable for image similarity. SSIM is a similarity measure between two images where for corresponding sub-windows of pixels, luminance, contrast, means, variances and covariances are evaluated. The mean of these calculations is taken to give the SSIM for the whole image. PSNR is a way of contextualizing and standardizing the MSE in terms of the pixel values of the image, with a higher PSNR corresponding to more similar images. PCC is used to measure the linear correlation of pixels in the images.

PSNR is normalized to the maximum potential pixel value, taken as 255 when the images are converted to 8-bit. For this dataset the PSNR appeared as high as the maximum 8-bit pixel values for each image was generally lower than the theoretical maximum value. Only SSIM can be interpreted as a fully-normalized metric, with values between 0 and 1 (1 being a perfect match).

### 2.14 Morphological Feature-Level Evaluation

Pairwise Spearman correlations [31] between the features in test set data were calculated for each model, with the mean values for each feature group grouped into correlation matrices, and visualized as heatmaps [19]. These features were split into several categories – area/shape, colocalization, granularity, intensity, neighbors, radial distribution and texture. We also visualized feature clustering using uniform manifold approximations (UMAPs) [32], implemented in python using the UMAP package [33].

### 2.15 Toxicity Prediction

We normalized the morphological profiles to zero mean and standard deviation of one and classified the compounds with K-NN-classifier (k = 5, Euclidian distance) into two groups using the controls as training label (positive controls were used as an example of toxic phenotype). To account for the unbalance between the number of positive and negative controls, we sampled with a replacement equal number of profiles from both categories for training and ran the classifier 100 times and used majority voting for the final classification.

## 3 Results

To systematically investigate the utility of the label-free prediction of Cell Painting from brightfield images, we conducted the evaluation on three separate levels: image-level, morphological feature-level, as well as a downstream analysis on the profile-level to identify toxic compounds. We evaluated the two models on the test set of 148 unique compounds, 26 positive control wells (mitoxantrone), and 77 negative control wells (DMSO) that represented two experimental batches and 12 plates (Table 1). Examples and descriptions of cells in these different wells are presented in supplementary Figure A.

### 3.1 Image-level Evaluation

The mean values of the image-level metrics for the U-Net/**cWGAN-GP** models respectively were SSIM: 0.42/**0.44**, PSNR: 51.7/**52.9**, MSE: 0.46/**0.45**, MAE: 0.41/**0.39**, and PCC: 0.84/**0.84**, with the superior model in bold. cWGAN-GP achieved on average superior image-level performance; however, the difference was always within the standard deviation (Table 2). Visual inspection of the model generated images with the ground truth revealed strong resemblance (Figures 3, 4, 1). Most notably, both models struggled in predicting the fine structures in AGP and Mito channels, this was also reflected as lower image-level performance for the channels. Interestingly, only the plasma membrane of AGP channel was successfully predicted, whereas the models were not able to reproduce the actin filaments and the Golgi-apparatus structures. Similarly with the Mito channel, the models performed well overall in predicting the mitochondrial structures, but we observed lack in small granularity and detail that was present in the ground truth. In addition, we observed that the blocking effect from inference time sliding window was visible in the generated images. Nevertheless, DNA, ER, and RNA channels were better predicted, of which RNA achieved the best image-level performance.

**Table 2:** Image metrics for each channel for the two models. The best performing model for each channel metric is highlighted in bold.

| Imaging Channel | Model | SSIM | PSNR | MSE | MAE | PCC |
| --- | --- | --- | --- | --- | --- | --- |
| DNA | U-Net | $0.38 \pm 0.13$ | $51.8 \pm 1.7$ | $0.45 \pm 0.12$ | $0.41 \pm 0.05$ | $0.90 \pm 0.06$ |
|  | **cWGAN-GP** | **$0.39 \pm 0.14$** | **$52.4 \pm 1.5$** | **$0.39 \pm 0.10$** | **$0.39 \pm 0.04$** | **$0.90 \pm 0.06$** |
| ER | U-Net | **$0.49 \pm 0.09$** | **$48.0 \pm 1.3$** | **$1.07 \pm 0.30$** | **$0.58 \pm 0.09$** | $0.86 \pm 0.06$ |
| | cWGAN-GP | $0.48 \pm 0.09$ | $47.8 \pm 1.3$ | $1.12 \pm 0.32$ | $0.61 \pm 0.09$ | **$0.87 \pm 0.06$** |
| RNA | U-Net | $0.59 \pm 0.07$ | $56.0 \pm 1.6$ | $0.18 \pm 0.09$ | $0.27 \pm 0.06$ | $0.92 \pm 0.06$ |
|  | **cWGAN-GP** | **$0.60 \pm 0.07$** | **$56.3 \pm 1.7$** | **$0.17 \pm 0.10$** | **$0.26 \pm 0.06$** | **$0.92 \pm 0.06$** |
| AGP | U-Net | $0.35 \pm 0.08$ | $52.4 \pm 1.8$ | $0.27 \pm 0.10$ | $0.39 \pm 0.06$ | $0.80 \pm 0.07$ |
|  | **cWGAN-GP** | **$0.37 \pm 0.09$** | **$54.7 \pm 1.3$** | **$0.23 \pm 0.09$** | **$0.34 \pm 0.06$** | **$0.82 \pm 0.06$** |
| Mito | U-Net | $0.31 \pm 0.06$ | $50.5 \pm 1.1$ | **$0.34 \pm 0.08$** | $0.42 \pm 0.05$ | **$0.74 \pm 0.08$** |
| | **cWGAN-GP** | **$0.35 \pm 0.06$** | **$53.1 \pm 1.4$** | $0.34 \pm 0.12$ | **$0.36 \pm 0.04$** | $0.70 \pm 0.08$ |

**Figure 3:**
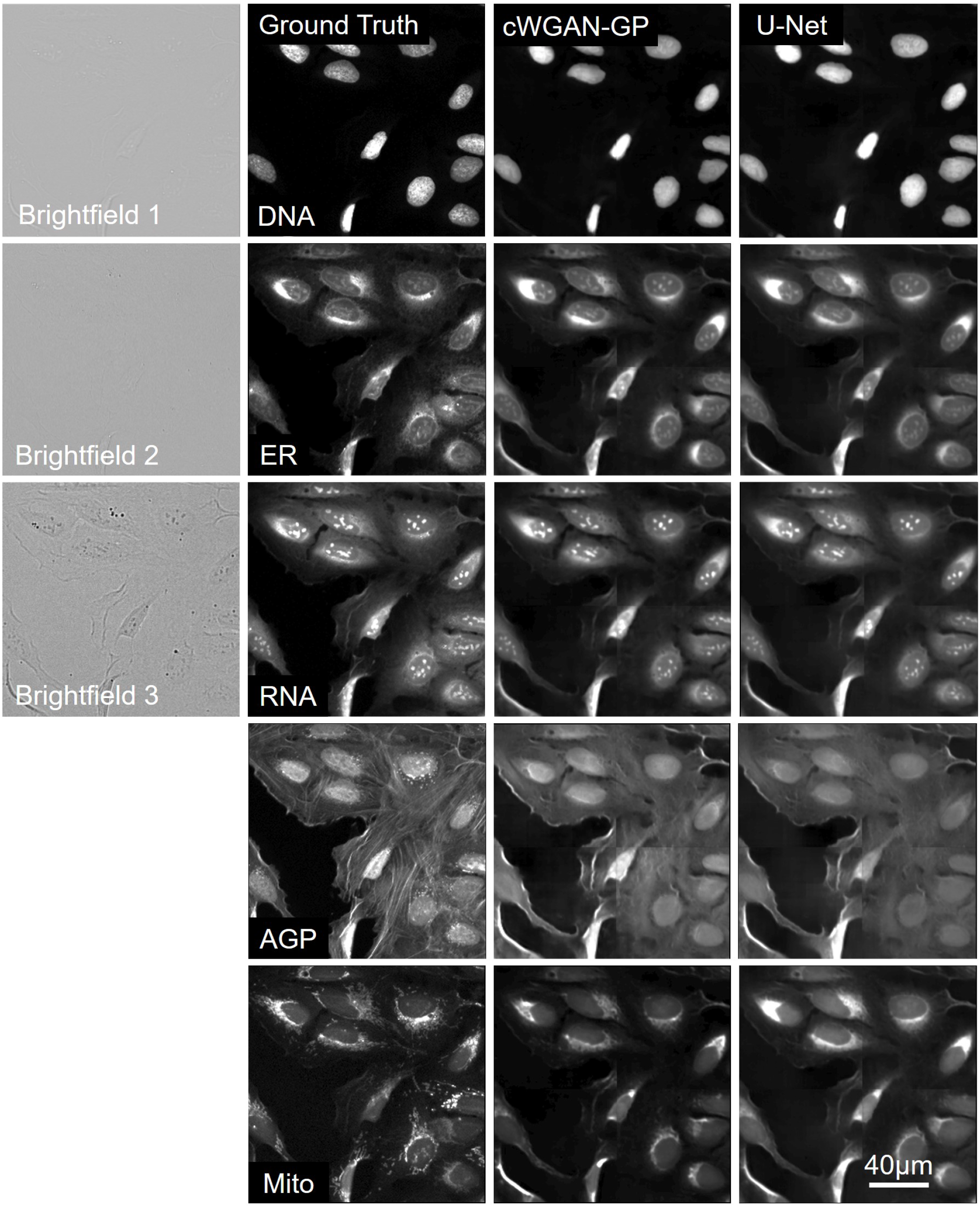
Cropped images of typical brightfield images, ground truth fluorescent, and predicted channels from the test dataset (Table 1) for the U-Net and cWGAN-GP models. Presented at scale of the input and output size used in the model networks (256 x 256 pixels), the first column contains the three brightfield z-planes which form the input, and the subsequent columns are the five fluorescent output channels. The stitching method has been applied to the model predicted images, and small artefacts of this process are visible. Images are independently contrasted for visualization.

**Figure 4:**
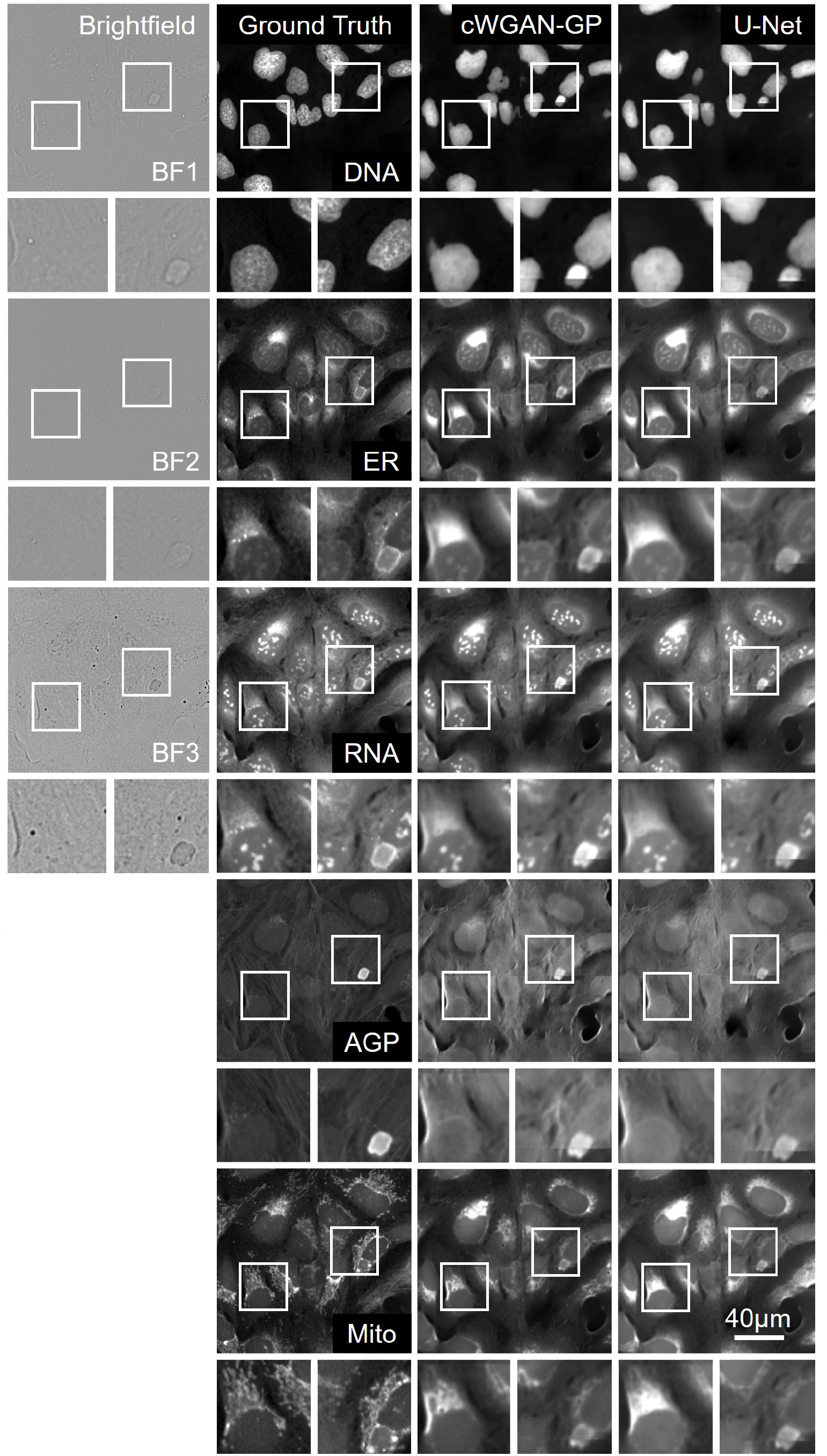
Two small outsets for each image show a magnified view of the selected region of the cell, and can be used to compare details between the ground truth and the two model outputs in each channel. Images are independently contrasted for visualization.

### 3.2 Morphological Feature-level Evaluation

Next, we extracted morphological features at the single-cell level and aggregated them to well-level profiles with the aim to further elucidate the utility of label-free Cell Painting. For consistency and to avoid any potential bias favoring the trained models, we chose to use the ground truth images for feature selection and reduced the profiles extracted from the predicted images accordingly. This resulted in 273 well-aggregated profiles consisting of 611 morphological features.

Using correlation analysis, we deconvoluted the profile similarities according to feature type, cell compartment, and imaging channels for both models (Figure 5). Overall, many morphological features extracted from the generated images showed substantial correlation with those extracted from ground truth images. Examples of accurately reproduced (>0.6 correlation) feature groups across both models were texture measurements of the AGP channel in both cells and cytoplasm; intensity measurements of the Mito channel in the cytoplasm, and granularity and texture measurements of the DNA channel within the nuclei. The highest performing feature group were granularity measurements of the RNA channel within the cytoplasm (0.86 correlation in both models). Almost all features correlated positively with the ground truth, and only a small number of features showed close to zero or negative correlation, such as the radial distribution of RNA in cytoplasm and the intensity of the AGP channel in nuclei. The cell colocalization features were not calculated by CellProfiler however the cytoplasm and nuclei colocalization features represent the cell as a whole.

**Figure 5:**
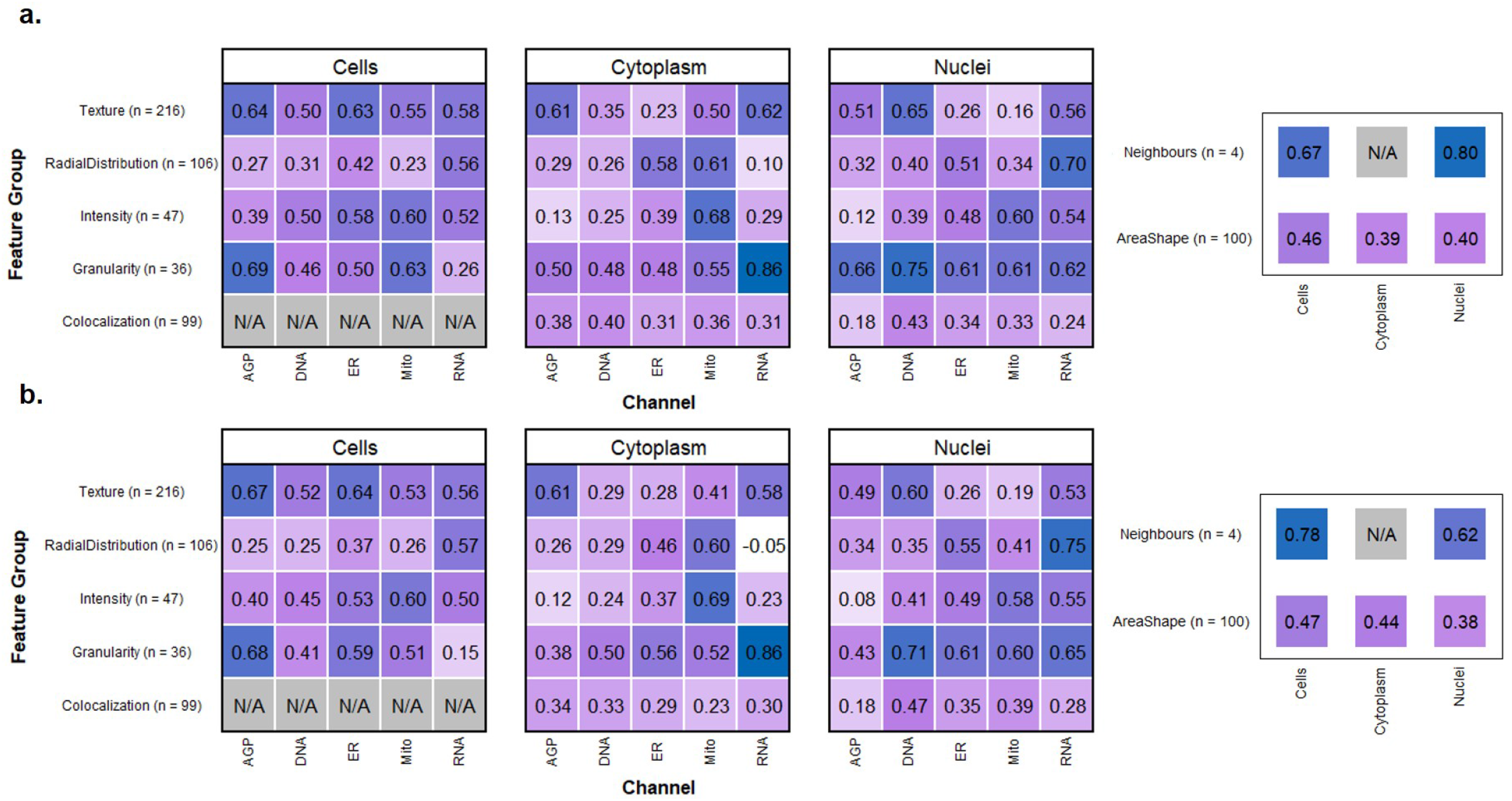
a systematic breakdown of correlation between morphological profiles. The features selected from the CellProfiler pipeline are presented as correlation heatmaps in 5.a (U-Net) and 5.b (cWGAN-GP). The correlations for each model are with the features extracted from the ground truth Cell Painting images, and are aggregated by channel and feature group for cell, cytoplasm, and nuclei objects. The number of features for each feature group is also presented. U-Net mean = 0.43, cWGAN-GP mean = 0.45.

The mean correlation of all the feature groups was 0.43 for the U-Net and 0.45 for cWGAN-GP, supporting the earlier finding of cWGAN-GP’s superiority in the image-level evaluation. The feature breakdown by cell group was: 169 cell features (mean correlation: 0.48/**0.50**), 217 nuclei features (**0.43**/0.41), and 225 cytoplasm features (0.38/**0.42**). The feature breakdown by feature type was, in descending order of best to worst mean correlation: 4 neighbors (0.70/**0.74**), 36 granularity (0.53/**0.55**), 216 texture (0.47/**0.49**), 47 intensity (0.46/**0.48**), 103 area/shape (0.42/**0.43**), 106 radial distribution (0.38/**0.39**) and 99 colocalization (0.31/**0.33**). By channel, the mean feature correlations were: RNA: 0.47/**0.49**, AGP: 0.43/**0.44**, DNA: 0.41/**0.44**, Mito: 0.40/**0.43**, ER: **0.42**/0.40.

The mean correlations for the top 10% of the selected features were 0.80/**0.81**. The mean correlations for the top 50% of the selected features were 0.64/**0.65**. The number of features with a correlation greater than 0.8 were 26/**30** (U-Net/**cWGAN-GP**), and for both models all feature groups and cell compartments had at least one feature with such a strong correlation, except for the colocalization feature group. For the cWGAN-CP model the breakdown of the 30 features with greater than 0.8 correlation was as follows: 12 cell, 11 nuclei, 7 cytoplasm, and this included the feature types: 11 texture, 7 radial distribution, 5 area/shape, 4 granularity, 2 texture, 1 neighbors, 0 colocalization.

Using uniform manifold approximation (UMAP), we visualized the high-dimensional morphological profiles for identification of underlying data structures (Figure 6). We observed that all three image sources (ground truth, U-Net, and cWGAN-GP) were separated in the common feature space due to the difference in the ground truth and predicted images. Despite this separation, we also observed that all the image sources maintained the overall data structure. The clear batch effect visible in the ground truth is also evident in the predicted images, and similarly the clustering of positive and negative control wells respectively is retained, indicating successful model performance. Features extracted from the cWGAN-GP model lie closer to the ground truth features than the U-Net extracted features, yet features from both models are closer to each other than either are to the ground truth.

**Figure 6:**
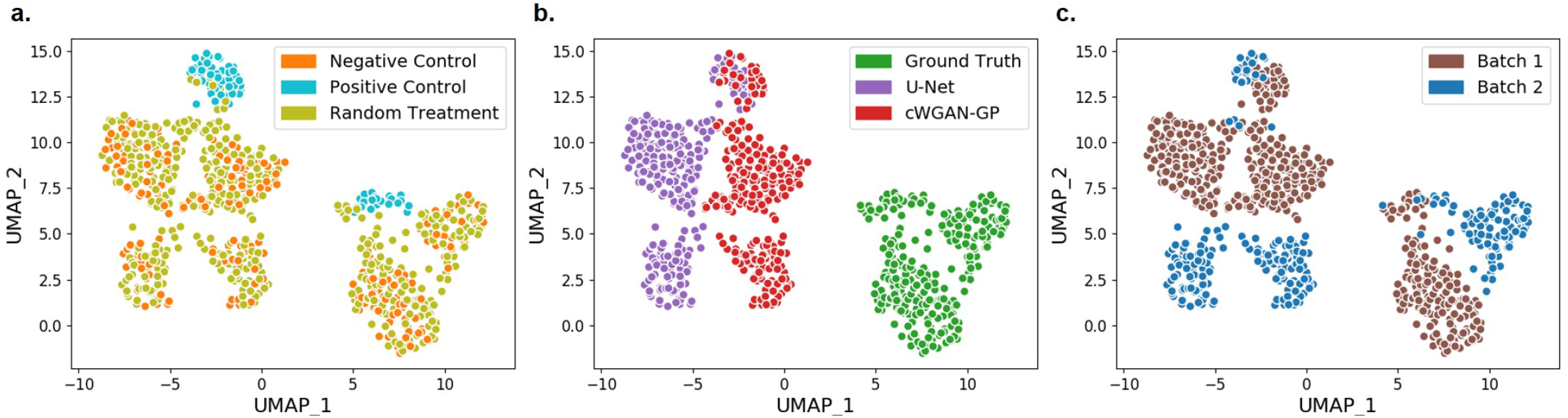
The UMAPs demonstrate that both models could reproduce the separation between treatments and batches seen in the ground truth features and that the clustering patterns are very similar. cWGAN-GP was closer to the ground truth than U-Net, although the two models sit much closer to each other in feature space than either to the ground truth.

### 3.3 Profile-Level Evaluation

Conclusions made from the label-free prediction of Cell Painting images should show agreement with ground truth datasets to be of experimental value. We therefore performed a series of analyses to identify compounds that elicit a comparable toxic phenotype to an established positive control compound, mitoxantrone. Our UMAP (Figure 6) showed that, as would be expected, most of the compounds showed greater resemblance to negative (DMSO) controls. Promisingly, some compounds clustered with our positive control compound. To identify these compounds, we trained a K-NN classifier using the control profiles as our training set. The classification resulted in identification of eight compounds in the ground truth, five in the cWGAN-GP model, and eight in the U-Net model profiles. The U-Net model achieved a sensitivity of 62.5% and specificity of 98.0% in toxicity classification whilst the respective values for cWGAN-GP were 50.0% and 99.3%.

## 4 Discussion

As the first full prediction of five-channel Cell Painting from brightfield input, we have presented evidence that label-free Cell Painting is a promising and practicable approach. We show our model can perform well in a typical downstream application, and that many label-free features from the predicted images correlate well with features from the stained images. In addition, we see success in clustering by treatment type from the features. We present indications of which channels and biomarkers can be satisfactorily by testing our model predictions with a comprehensive segmentation-based feature extraction profiling methodology.

Our results demonstrate that incorporating adversarial loss results in a small increase in performance over L1 loss based on the pixel-wise evaluation metrics in all channels except ER (Table 2). In domain transfer problems such as ours, the finer details in some of the predicted channels can be obstructed and blurred with a pixel-wise loss function such as L1 loss [34]. Just a small performance difference is expected as the network and loss function for training both models were very similar, and in the cWGAN-GP model the adversarial component of loss is weighted relatively low compared to L1. In addition, features from images from the cWGAN-GP model show an increase in performance on the U-Net model, with slightly higher mean correlations to the ground truth. The strong correlation of specific feature groups increases confidence that extracted morphological feature data from predicted images can be used to contribute overall morphological profiles of perturbations.

Biologically, it is expected that correlations of feature groups within the DNA channel are higher within the nuclei compartment than either cytoplasm or cells since the nuclei compartment is morphologically very distinct from the cytoplasmic region. The high correlation of radial distribution features in the RNA channel suggests that successful visualization of the nucleoli within the nuclear compartment has a large effect on this particular feature group. AGP and Mito channels both contain small and subtle cellular substructures which are typically less than two pixels wide. We postulate that this fine scale information is not present in the brightfield image in our data, making accurate image reproduction of the AGP and Mito channels very challenging regardless of model choice. There are known limitations of brightfield imaging which will always restrict a domain transfer model with brightfield input. For example, brightfield images can display heterogeneous intensity levels and poor contrast. Segmentation algorithms have been proven to perform poorly when compared with fluorescence channels, even after illumination correction methods have been applied [27].

As shown in Figure 6, in UMAP feature space the model-predicted features do not overlap with the ground truth features. This highlights limitations of the model but also the batch effect – it is not expected for the model to predict an unseen batch. The relevant structure of the data is maintained although not absolute values. This is also notable at the image level, for example the high MSE is likely due to the batch effect causing a systematic difference in pixel values between the training and test batches. Similarity metrics such as SSIM may be more informative in this instance. It is notable that ground truth features from different batches also sit in non-overlapping feature space in our UMAPs. Within batches, the negative controls and treatments would form sub-clusters depending on the treatments in a larger dataset, however we acknowledge our test set was relatively small, resulting in minimal sub-clustering.

Other studies [21] which have used U-Net based models to predict fluorescence from transmitted light or brightfield have evaluated their performance on pixel-level metrics such as PCC [15], SSIM [28] [29] and PSNR [35] [24]. The mean PCC of all channels in our test set is 0.85 (using the best model for each channel), a value which compares favorably with prevailing work in fluorescent staining prediction [8]. In our data two channels (DNA, RNA) exceed a PCC of 0.90 for both models. However, absolute values of image metrics are heavily data dependent so we present these metrics primarily for model comparison.

Such metrics are standard in image analysis but have some limitations for cellular data. Treating each pixel in the image equally is a significant limitation as pixels representing the cellular structures are clearly more important than background (void) pixels [35]. Some channels such as the DNA channel are more sparse than other channels such as AGP, and as such the number of pixels of interest vs background pixels is different. Extracting features from scientific pipelines provides a more biologically relevant and objective evaluation to give deep learning methods more credibility and a greater chance of being practically employed in this field [36].

Despite the small sample size, the models performed well at predicting the toxic phenotype, with specificities of 98.0% (U-Net) and 99.3% (cWGAN-GP). The sensitives were 62.5% and 50% respectively. Although comparative or superior results could likely be achieved with an image classifier trained on the brightfield, as a label-free and image-to-image approach there is a lot of promise as a generalist model to perform multiple tasks. The simple visualization of Cell Painting channels is one approach to improve the interpretation of brightfield images. Our results highlight the rich information captured in the brightfield modality, which may currently be under-utilized in morphological profiling.

Our approach by itself cannot replace full fluorescent staining, but we have provided evidence that it may be possible to replicate the information of some Cell Painting channels and feature groups, and that the brightfield modality by itself may be sufficient for certain experimental applications. Importantly, employing such methods may reduce time, experimental cost and enable the utilization of specific imaging channels for experiment-specific, non-generic fluorescent stains. We acknowledge that particular feature groups which predict poorly in our models (such as colocalization features) may result in an inability (or reduced sensitivity) to identify cellular phenotypes which are characterized fully by these features. In such situations, the replacement of generic stains for phenotype- or target-relevant biomarkers may offer an effective solution to the standard Cell Painting protocol.

One limitation of this study is the dataset used. Future studies will require larger datasets with greater diversity in terms of compound treatments and collection sources. Matching the numbers of fields in our training and test sets to the size of a typical dataset used in drug discovery would allow for greater insight into the capabilities of label-free Cell Painting. International collaborations such as the Joint Undertaking of Morphological Profiling using Cell Painting (JUMP-CP) aim to further develop Cell Painting and provide a highly valuable public dataset for this use [37]. It is notable that we have evaluated predictions with downstream features the networks have not seen – simply extracted classically from the images. In future studies, incorporating feature information in the training of the network itself may drastically improve performance, and this task may be appropriate for a transfer learning approach [38].

In summary, we propose a deep learning approach which can digitally predict Cell Painting channels without the application of fluorescent stains, using only the brightfield imaging modality as an input. Building upon previous work [8] [22] we have predicted the five fluorescent channels and used these images to predict the associated groups of morphological features from a standard image analysis pipeline. We then used the features from the predicted images to assess how information-rich such images are. Finally we have provided a critical evaluation of the predictions using morphological features extracted from images using CellProfiler analysis and the resulting compound profiles.

## Acknowledgments

The authors would like to thank the AZ UKCCB team for the provision of the cell line. JCZ is funded by the UKRI-BBSRC DTP (UK Research and Innovation and the Biotechnology and Biological Science Research Council Doctoral Training Partnership) studentship and AstraZeneca. CBS acknowledges support from the Philip Leverhulme Prize, the Royal Society Wolfson Fellowship, the EPSRC grants EP/S026045/1 and EP/T003553/1, EP/N014588/1, EP/T017961/1, the Wellcome Innovator Award RG98755, the Leverhulme Trust project Unveiling the invisible, the European Union Horizon 2020 research and innovation programme under the Marie Skodowska-Curie grant agreement No. 777826 NoMADS, the Cantab Capital Institute for the Mathematics of Information and the Alan Turing Institute. Authors EM, GW, RT, YW are employees of AstraZeneca. AstraZeneca provided the funding for this research and provided support in the form of salaries for the authors, but did not have any additional role in the study design, data collection and analysis, decision to publish, or preparation of the manuscript.

## Data Availability Statement

The datasets generated and analyzed in this study are not publicly available due to AstraZeneca licenses but are available from the corresponding author on reasonable request. Source code from this project is available on GitHub: https://github.com/crosszamirski/Label-free-prediction-of-Cell-Painting-from-brightfield-images

1 https://github.com/crosszamirski/Label-free-prediction-of-Cell-Painting-from-brightfield-images

