## Supplemental Table 1 for "Label-Free Prediction of Cell Painting from Brightfield Images"

| **Imaging Channel** | **Stain** | **Stock Preparation** | **Stock Concentration** | **Dilution Factor** | **Working Concentration** |
| --- | --- | --- | --- | --- | --- |
| **DNA** | Hoechst 33342 (Thermo #H3570) | Use as provided | 10 mg/mL | 1:2000 | 5 µg/mL |
| **ER** | Concanavalin A / Alexa Fluor 488 (Thermo #C11252) | Add 1 mL 0.1M sodium bicarbonate (in dH_2_O) to vial | 5 mg/mL | 1:500 | 10 µg/mL |
| **RNA** | SYTO 14 green fluorescent nucleic acid stain (Thermo #S7576) | Use as provided | 5 mM | 1:555.5 | 9 µM |
| **AGP** | Wheat-germ agglutinin / Alexa Fluor 555  (Thermo #W32464) | Add 5 mL dH2O to vial, centrifuge at 10’000g for 30s to remove aggregates | 1 mg/mL | 1:666.7 | 1.5 µg/mL |
|  | Phalloidin / Alexa Fluor 568 (Thermo #A12380) | Add 1.5 mL 100% (v/v) methanol to vial | 1 mL/mL  (300 units in vial) | 1:200 | 5 µL/mL  (1.5 units) |
| **Mito** | MitoTracker Deep Red  (Thermo #M22426) | Add 91 µL DMSO to vial | 1 mM | 1:2000 | 0.5 µM |
