## Supplementary figures and images for "Label-Free Prediction of Cell Painting from Brightfield Images"

### Supplemental Figure 1

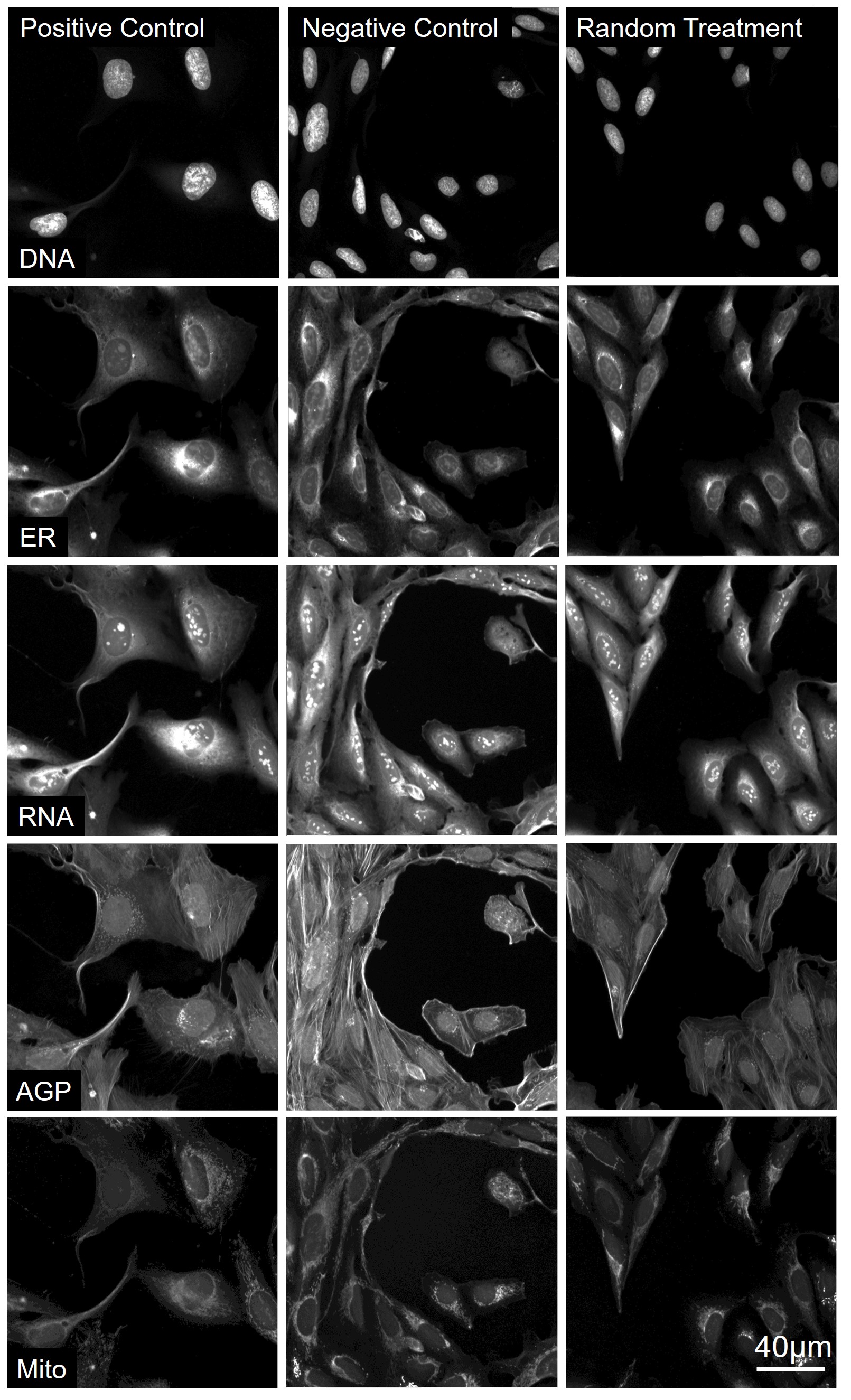

### Supplemental Figure 2

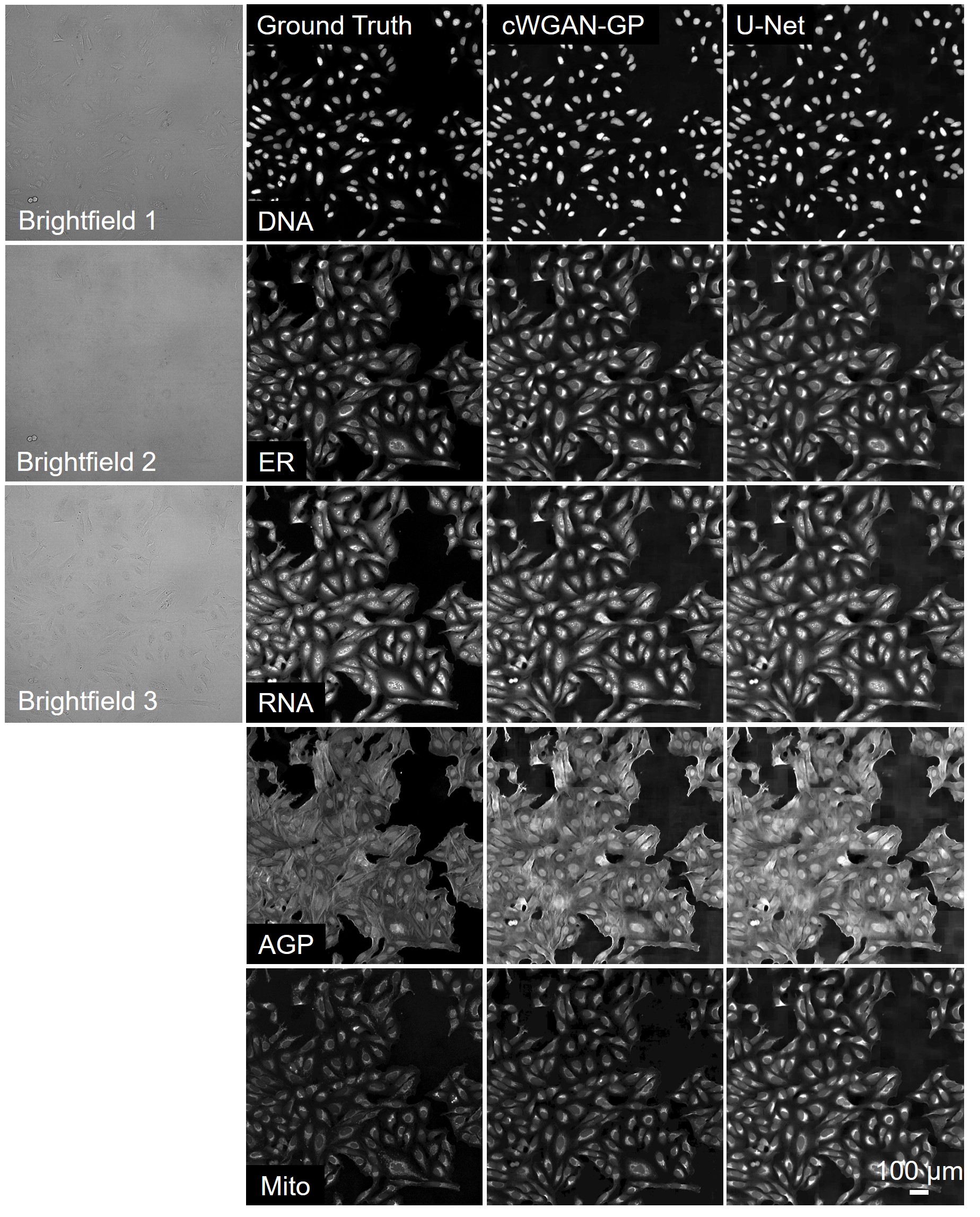
